# Spatio-temporal machine learning for multi-horizon prediction of bluetongue outbreaks

**DOI:** 10.64898/2026.05.21.726753

**Authors:** Lucy M. Devlin, Phi H Nguyen, Ross N. Cuthbert, Phu Ngoc Doan, Viet Hung Tran, Zichi Zhang, Archie K. Murchie, Connor G. G. Bamford, Jaimie T. A. Dick, Eric R. Morgan, Thai S. Mai

## Abstract

Reliable early warning of infectious disease outbreaks remains a major challenge for surveillance systems, particularly for vector-borne pathogens whose transmission depends on interactions among hosts, vectors, and climate-sensitive environmental conditions. Data-driven forecasting offers a promising approach for predicting outbreak risk using surveillance and environmental data. This study develops a logit-weighted ensemble (LWE), a machine-learning framework that predicts outbreak occurrence 1–6 months ahead at the administrative unit–month scale using routinely available outbreak notifications and gridded climate data. Bluetongue virus (BTV), an arbovirus of ruminants transmitted by *Culicoides* biting midges, provides a well-characterised system in which transmission is strongly shaped by climate, making it a useful system for applying and testing this approach. The framework is evaluated using surveillance data collected between 2005 and 2024 from France, Greece, and Italy, selected for their long-running and high-quality outbreak surveillance records. Across all three countries, the LWE achieved the strongest and most stable predictive performance under a recall-focused evaluation that prioritises correctly identifying outbreak months. It outperformed or matched 14 benchmark models, with differences becoming more pronounced at longer lead times (month +3 onward), when predictions are more uncertain and outbreaks are relatively rare. Predictability varied across countries, with the highest performance in Greece, strong performance in France, and lower, more variable performance in Italy, reflecting differences in how consistently outbreaks occur and spread across regions. Overall, the results demonstrate that horizon-aware, climate-informed forecasting can reliably identify months and locations at elevated risk of outbreak occurrence up to six months in advance, supporting surveillance planning and preparedness across heterogeneous European settings. The ensemble framework provides a robust and portable strategy for outbreak prediction using routinely collected surveillance and environmental data.

**Author Summary:** Predicting infectious disease outbreaks before they occur remains a major challenge, particularly for diseases influenced by environmental conditions. In this study, we focus on bluetongue, a viral disease of livestock transmitted by biting midges, where transmission is strongly affected by climate and seasonal patterns. We develop a method that uses routinely collected outbreak reports and climate data to estimate where and when outbreaks are more likely to occur, up to six months in advance. We apply this approach across three European countries with a history of bluetongue outbreaks. We find that combining climate information with recent outbreak patterns can provide useful early signals of increased risk. Predictions are most accurate at shorter timeframes, but longer-range forecasts can still support planning and preparedness. Because our approach uses widely available data, it could be applied in other regions or to similar environmentally driven diseases. However, it does not include factors such as vaccination, animal movement, or detailed information on vector populations, which may also influence how outbreaks develop.

**Graphical Abstract:** 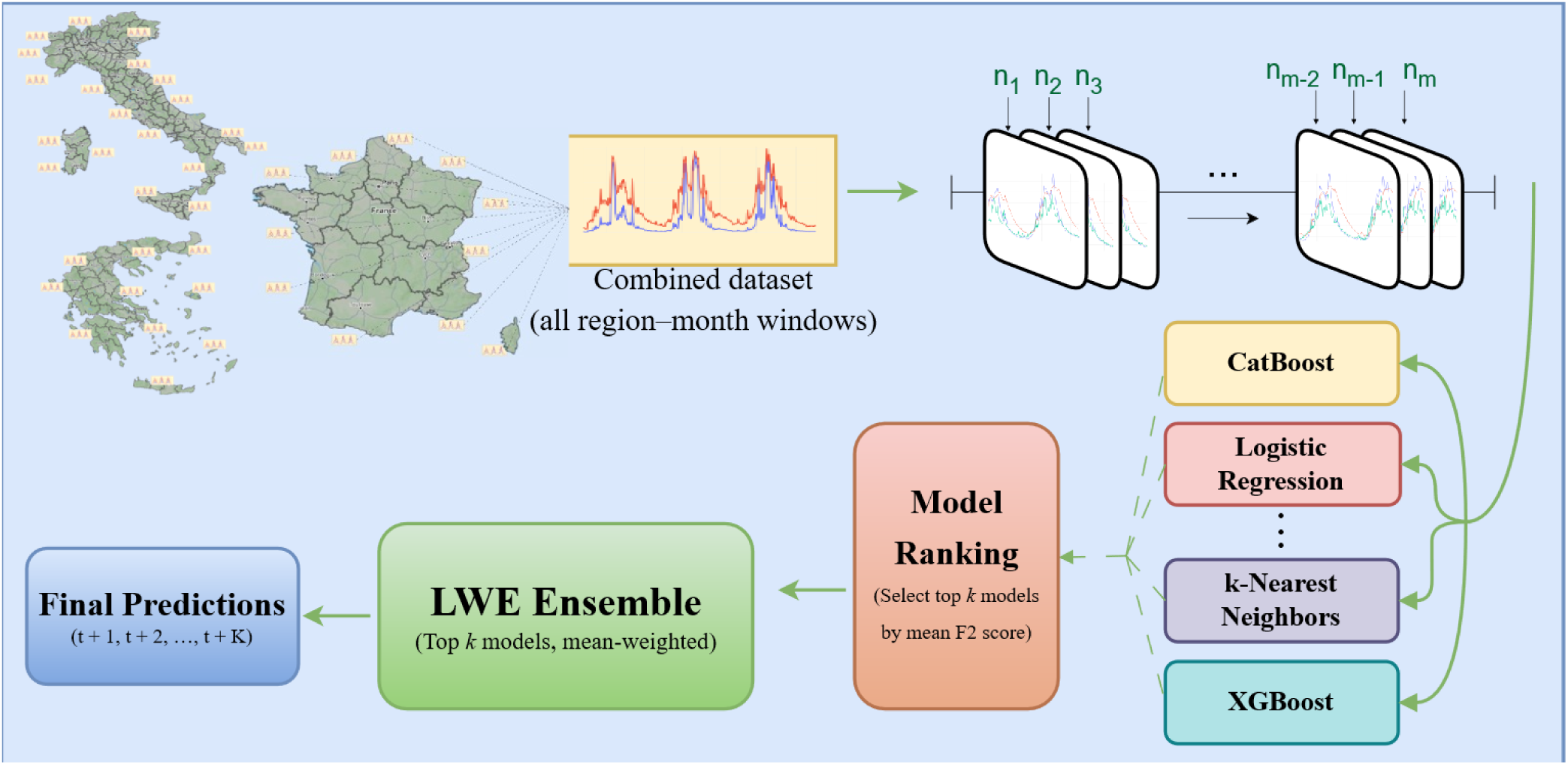

## 1. Introduction

Accurately predicting infectious disease outbreaks before they occur remains a central challenge in epidemiology and animal health (1, 2). Despite advances in statistical and mechanistic modelling, reliable early warning is difficult because transmission processes are nonlinear, spatially heterogeneous, and shaped by interacting environmental and surveillance dynamics (3, 4). Forecasting systems must therefore operate under substantial uncertainty and severe class imbalance, where the imbalance arises from the relatively small number of outbreak occurrences compared with non-outbreak observations at the level of individual locations and months, while still producing actionable outputs for real-world surveillance and preparedness (5).

These challenges are particularly pronounced for vector-borne diseases (VBDs), where transmission depends on interactions among hosts, pathogens, vectors, and the environment. Because vector survival, activity, and pathogen replication are climate-sensitive, transmission is shaped by temperature and seasonality, which regulate vector abundance, biting rates, and the extrinsic incubation period (6–8). As a result, outbreak risk is highly seasonal, while climate change is reshaping vector distributions, extending transmission periods, and enabling spread into previously unaffected temperate regions (9, 10). Evidence of cold tolerance and overwintering in some vector populations suggests potential for local persistence between seasons, while anthropogenic changes such as irrigation, land-use change, and intensive livestock production may further enhance breeding opportunities (11–14).

Bluetongue disease exemplifies these challenges as a globally important viral infection of domestic and wild ruminants transmitted by *Culicoides* biting midges (15–17). Vertical transmission has also been reported (18–20). Clinical signs range from fever, oedema, and oral ulceration to reproductive losses and reduced productivity, contributing to substantial economic impact. Endemic across most continents except Antarctica (14), BTV remains a major concern for livestock production and international trade (21–24). The resulting economic burden includes direct production losses and control-related costs, including vaccination, diagnostics, vector control, surveillance, and outbreak response (25–27). Control is further complicated by multiple circulating serotypes, which hinder effective vaccination and population immunity (14, 15, 18, 25), while response effectiveness is constrained by delayed detection, high resource demands, and limited predictive capacity (18, 28). In practice, control measures such as vaccination are deployed reactively after outbreak detection, concentrating resources in affected areas and allowing continued spread into neighbouring susceptible populations, with surveillance signals arising only after transmission is established and limiting lead time for intervention (29, 30).

Predictive approaches enable forward-looking spatiotemporal risk assessment, supporting early warning and risk stratification at administrative scales (31, 32). In the context of climatic variability and expanding vector suitability, climate-informed prediction is central to developing prospective BTV risk models under shifting environmental conditions (7, 9). Mechanistic and vector-based models have long been used to describe BTV transmission dynamics and to explore climate-driven suitability, offering biological interpretability and support for scenario analysis (33, 34). More recently, forecasting efforts have increasingly incorporated data-driven and machine-learning approaches, including tree-based and neural network models, to predict regional outbreak risk using routinely collected surveillance and environmental data, sometimes combined with atmospheric dispersion modelling to assess windborne incursion (33, 35, 36). Existing studies are, however, often limited to single settings or short-term forecasts and rely on retrospective evaluations misaligned with early warning under severe class imbalance. As a result, evidence on the robustness and operational stability of multi-horizon outbreak forecasts, defined as predictions made several months in advance, remains limited.

This study develops and evaluates a multi-horizon machine-learning framework to predict bluetongue outbreak occurrence at the unit–month administrative scale using surveillance data from France, Greece, and Italy, which have recurrent BTV outbreaks and established surveillance systems. Here, forecast horizon refers to the months between prediction and the target outbreak period. Using climate summaries and recent outbreak history, probabilistic forecasts 1–6 months ahead are generated and evaluated under strict temporal validation, with models trained on past data and tested on future observations. Multiple classifier families are benchmarked and combined within a decision-aligned ensemble, termed the Logit-Weighted Ensemble (LWE), which integrates selected base learners to improve robustness and reduce overconfident predictions under severe class imbalance (37).

The specific aims are to (i) produce probabilistic outbreak predictions at lead times of 1-6 months; (ii) evaluate performance within a time-respecting, operationally realistic framework; (iii) quantify the contribution of the LWE under severe class imbalance; and (iv) examine how predictability, model ranking, and performance variability change with increasing forecast horizon across heterogeneous reporting systems. Collectively, these aims evaluate the feasibility and stability of multi-horizon BTV forecasting under real-world surveillance constraints and demonstrate a framework transferable to other climate-sensitive diseases using routine environmental data for prospective risk prediction.

## 2. Results

### 2.1 Pooled benchmark performance across forecast horizons

Pooled predictive performance across forecast horizons shows two consistent patterns across benchmark model families. First, performance decreases monotonically with increasing lead time for all models, reflecting the expected loss of predictive signal as the forecast target moves further from the most recent outbreak history and climate observations. This pattern is consistent with the epidemiology of BTV, where recent transmission and short-term climatic conditions influencing *Culicoides* vector activity provide stronger signals of ongoing outbreak risk than longer-term forecasts. Second, the strongest benchmark performance is dominated by tree-based ensembles, particularly gradient-boosted methods. Among individual models, XGBoost achieves the highest mean pooled *𝐹*_2_ score (0.822), followed closely by LightGBM (0.816) and a group of competitive tree-based baselines, including Balanced Random Forest, HistGradientBoosting, and Gradient Boosting. The small absolute separation among these top-performing models indicates that multiple high-capacity tree learners extract broadly similar signal from the unit–month feature representation.

### 2.2 Dependence of model performance on forecast horizon

Model rankings are not constant across forecast horizons. At short lead times (Month +1 and Month +2), several tree-based ensembles attain very high pooled performance (typically F₂ ≥ 0.90), with minimal separation among the top-ranked methods (Figure 1). This indicates that most models capture similar short-term persistence signals present in the data, likely reflecting ongoing local transmission and favourable vector activity within the same seasonal transmission period. From Month +3 onward, as predictive signal weakens, separation among models becomes more pronounced (Figure 1B). This reduction in predictive signal is consistent with the increasing uncertainty in vector abundance, host exposure, and local transmission conditions over longer forecast horizons. Boosted tree methods remain consistently competitive at longer horizons, whereas some model classes exhibit steeper performance degradation. This pattern is most evident for margin-and kernel-based support vector machine variants, which show reduced robustness under horizon-dependent score shifts and severe class imbalance.

**Figure 1:**
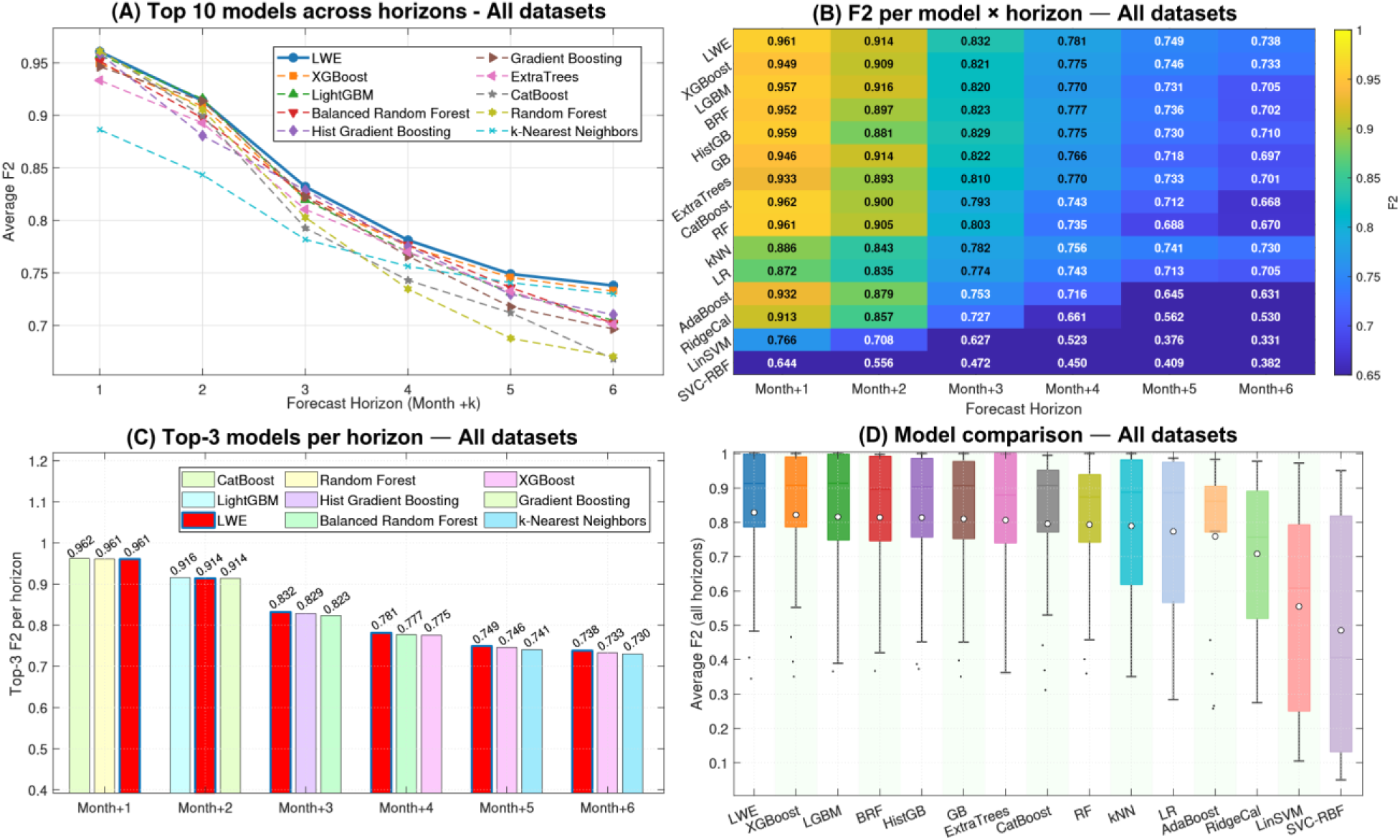
Pooled multi-horizon forecasting performance across France, Italy, and Greece (micro-averaged). Performance of all benchmark models and the logit-weighted ensemble (LWE) across six forecast horizons (Month +1 to Month +6), evaluated on the held-out test period and pooled across all unit–month windows. **(A)** Mean *𝐹*_2_ score by forecast horizon for the top-performing models, illustrating the expected decline in performance with increasing lead time. **(B)** Heatmap of *𝐹*_2_ scores by model and horizon, highlighting horizon-dependent changes in relative model ranking. **(C)** Horizon-wise top three models ranked by test *𝐹*_2_; small ranking changes at short horizons reflect minor absolute score differences and are not interpreted as substantively meaningful. **(D)** Distribution of *𝐹*_2_ scores across evaluation slices (unit–horizon combinations), summarising performance variability across space and time; tighter distributions indicate greater stability under heterogeneous surveillance conditions. Together, the panels show that while multiple models perform similarly at short horizons, performance diverges as lead time increases, with the LWE providing the most stable performance at longer horizons.

### 2.3 Recall–precision trade-offs under the 𝐹_2_ objective

Performance differences among benchmark models are further clarified by examining recall–precision trade-offs at the horizon-specific decision thresholds (Table 1). These trade-offs are particularly relevant for infectious disease surveillance, where failing to detect an outbreak month may delay control measures such as vaccination or movement restrictions, potentially allowing further local transmission. Among the strongest-performing models, XGBoost operates at a relatively balanced operating point, achieving a mean recall of 0.835 and a mean precision of 0.784. In contrast, Gradient Boosting and Random Forest achieve the highest mean precision values (0.825 and 0.823, respectively), albeit at slightly lower recall. Models that prioritise recall more aggressively show distinct trade-offs. Logistic Regression attains the highest mean recall (0.893) but with substantially lower precision (0.630), indicating frequent positive classifications under the recall-weighted objective. Balanced Random Forest exhibits a similar recall-forward profile (recall 0.866; precision 0.699), consistent with its class-balancing design.

**Table 1:**
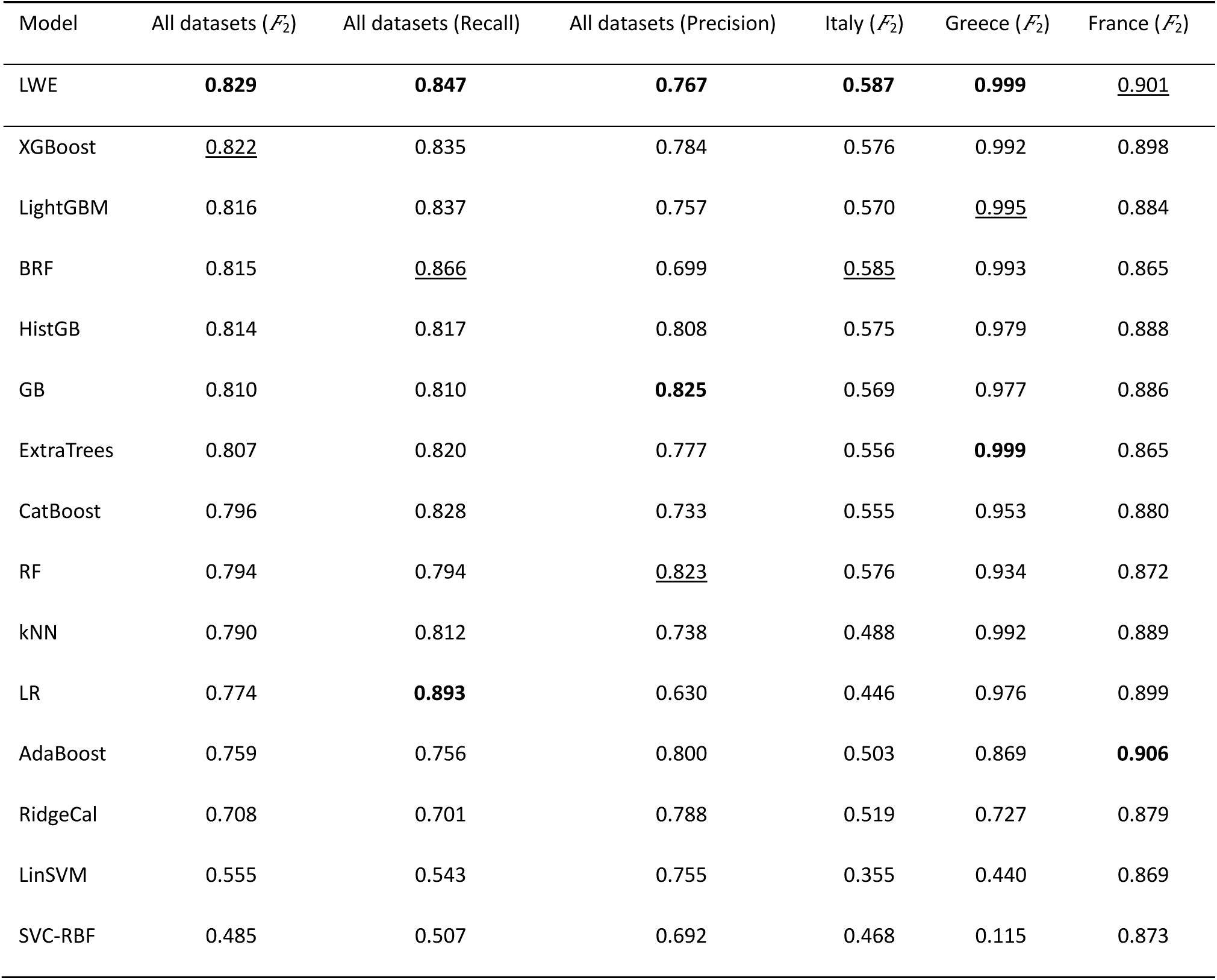
Model comparison under the primary objective (*𝐹*_2_). Mean *𝐹*_2_ is micro-averaged over all unit–month test windows and then averaged across horizons (Month +1 to Month +6). For the pooled setting (All datasets), we additionally report mean recall and mean precision, computed at the horizon-specific thresholds selected on validation, to summarise the implied operating point. Columns for Italy, Greece, and France report mean *𝐹*_2_ only. For transparency, all metrics are computed on the held-out 2022–2024 test period. Bold indicates the best value in each column and underlined indicates the second-best value (ties allowed).

| Model | All datasets ($\mathcal{F}_2$) | All datasets (Recall) | All datasets (Precision) | Italy ($\mathcal{F}_2$) | Greece ($\mathcal{F}_2$) | France ($\mathcal{F}_2$) |
| --- | --- | --- | --- | --- | --- | --- |
| LWE | **0.829** | **0.847** | **0.767** | **0.587** | **0.999** | <u>0.901</u> |
| XGBoost | <u>0.822</u> | 0.835 | 0.784 | 0.576 | 0.992 | 0.898 |
| LightGBM | 0.816 | 0.837 | 0.757 | 0.570 | <u>0.995</u> | 0.884 |
| BRF | 0.815 | <u>0.866</u> | 0.699 | <u>0.585</u> | 0.993 | 0.865 |
| HistGB | 0.814 | 0.817 | 0.808 | 0.575 | 0.979 | 0.888 |
| GB | 0.810 | 0.810 | **0.825** | 0.569 | 0.977 | 0.886 |
| ExtraTrees | 0.807 | 0.820 | 0.777 | 0.556 | **0.999** | 0.865 |
| CatBoost | 0.796 | 0.828 | 0.733 | 0.555 | 0.953 | 0.880 |
| RF | 0.794 | 0.794 | <u>0.823</u> | 0.576 | 0.934 | 0.872 |
| kNN | 0.790 | 0.812 | 0.738 | 0.488 | 0.992 | 0.889 |
| LR | 0.774 | **0.893** | 0.630 | 0.446 | 0.976 | 0.899 |
| AdaBoost | 0.759 | 0.756 | 0.800 | 0.503 | 0.869 | **0.906** |
| RidgeCal | 0.708 | 0.701 | 0.788 | 0.519 | 0.727 | 0.879 |
| LinSVM | 0.555 | 0.543 | 0.755 | 0.355 | 0.440 | 0.869 |
| SVC-RBF | 0.485 | 0.507 | 0.692 | 0.468 | 0.115 | 0.873 |

### 2.4 Stability of performance across units and horizons

The consistency of model performance across unit–horizon slices is illustrated in Figure 1D. Leading tree-based ensembles achieve high *𝐹*_2_ scores with relatively limited spread, indicating stable performance across administrative units and forecast horizons. This suggests that the models capture predictive signals that remain informative across heterogeneous transmission environments, where climatic conditions, vector activity, and livestock distributions vary across regions. In contrast, weaker model families exhibit both lower average performance and substantially greater variability. This pattern is most pronounced for support vector machine (SVM) variants, which show wide dispersion in scores across units and horizons. Overall, these results indicate that boosted and bagged tree ensembles provide not only strong average performance but also greater stability for multi-horizon BTV forecasting in this setting.

## 3. Discussion

### 3.1 Key findings and interpretation

This study shows that BTV outbreak occurrence at the unit–month reporting scale can be forecasted 1-6 months ahead using routinely available climate summaries together with previous outbreak data, highlighting the potential of data driven forecasting to support early warning for climate sensitive infectious diseases. Across three epidemiologically and operationally distinct settings (France, Greece, and Italy), these inputs consistently supported prospective early warning of periods and locations with higher outbreak likelihood, demonstrating that meaningful predictive signal is present even when models rely only on information that is routinely collected and updated within surveillance systems.

As expected, predictive performance declined with increasing lead time (i.e., the interval between when a forecast is generated and the month for which outbreak risk is predicted), reflecting growing uncertainty as forecasts move further from the most recent outbreak observations and meteorological conditions. Importantly, however, skill remained operationally informative through Month +6 under a recall-oriented objective, which prioritises correctly identifying outbreak months even at the cost of some additional false alerts. This indicates that BTV risk at the reporting-unit scale is shaped not only by short-term persistence but also by slower seasonal and environmental processes that provide advance warning beyond the immediate near term. Across both pooled and country-specific evaluations, the strongest-performing methods were consistently nonlinear, particularly tree-based ensemble learners. This pattern indicates that the relationship between climate conditions, recent outbreak history, and reported outbreak occurrence is structured but not well represented by simple linear effects. Because the outcome variable is derived from routine surveillance notifications rather than true infection prevalence, model performance necessarily reflects a combination of underlying transmission dynamics and properties of the reporting process. Nevertheless, the fact that similar model families ranked highly across countries with markedly different prevalence levels, administrative granularity, and outbreak histories suggests that the learned signal is not purely an artefact of reporting practice but captures reproducible structure relevant to outbreak occurrence at the surveillance scale. These countries also span contrasting epidemiological contexts, suggesting that the model rankings are robust across differing transmission regimes.

Among all evaluated approaches, the logit-weighted ensemble (LWE) exhibited the most stable behaviour as forecast horizon increased. While several high-capacity single learners performed comparably at short lead times, differences between models became more pronounced from Month +3 onward, when class imbalance, calibration drift, and horizon-specific shifts in score distributions increasingly influenced performance. As these horizon-related effects became more pronounced, the LWE consistently matched or exceeded the best individual models, supporting the interpretation that combining complementary decision functions reduces sensitivity to horizon-dependent instability and yields more robust early warning signals under heterogeneous conditions.

### 3.2 Climate-driven constraints on BTV transmission

The strong and consistent predictive performance obtained using only monthly climate variables and recent outbreak history indicates that climate exerts a dominant and coherent influence on BTV outbreak occurrence at the administrative unit–month scale. Across all three countries, skill remained high even as forecasts extended several months ahead, suggesting that climate is not merely capturing short-term persistence, but is encoding broader seasonal and environmental constraints on transmission. This is consistent with the well-established sensitivity of *Culicoides* vectors to temperature and moisture, whereby relatively small changes in thermal and hydrological conditions can strongly affect vector survival, activity, biting rates, and the speed of viral replication within the vector (6, 7, 38). Because these processes operate on seasonal rather than daily timescales, coarse monthly summaries are sufficient to capture much of the biologically relevant variation that determines whether transmission can be sustained within a given reporting unit. Beyond this, the results suggest that climate provides advance information on outbreak risk beyond the immediate near term. Increasing evidence that winter and early and late season temperatures are becoming more variable, including more frequent warm spells, provides a plausible mechanism by which climate conditions outside the main transmission period can influence subsequent outbreak risk by altering phenology and early-season amplification of vector populations (7, 38). The fact that similar climate predictors supported forecasting across countries with different surveillance systems and outbreak histories further suggests that the models are learning generalisable ecological constraints on transmission, rather than exploiting context-specific reporting patterns. In this context, short outbreak-history lags act as a complementary signal that captures ongoing transmission and delayed detection, while climate variables encode the broader environmental envelope within which BTV emergence and persistence are possible.

### 3.3 Horizon dependence and practical lead-time trade-offs

The usefulness of outbreak forecasts depends strongly on lead time, not only because predictive uncertainty increases with horizon, but because the appropriate interpretation of model outputs changes. At longer lead times of five to six months, predictions should be interpreted differently. Rather than serving as direct triggers for intervention, these forecasts are more appropriately viewed as early indicators of elevated risk that can inform preparedness, situational awareness, and resource planning, and that are subsequently refined as new data become available. This distinction highlights that forecast utility does not fully depreciate at longer horizons but instead shifts in how outputs should be operationally interpreted. In many outbreak responses, control resources are deployed primarily in locations where transmission has already been detected rather than in areas at imminent risk. When vaccine supply or field capacity is limited, this reactive allocation can result in interventions lagging behind the advancing transmission wave (39). Forecasts that identify elevated risk several months in advance may therefore help shift responses from reactive containment toward more anticipatory deployment of surveillance and vaccination resources.

Forecasts made one to two months ahead benefit strongly from recent outbreak history and short-term stability in meteorological conditions, supporting relatively high-confidence predictions. At intermediate lead times of three to four months, recent persistence contributes less and differences between modelling approaches become more apparent. In this setting, ensemble approaches such as the LWE provide particularly stable guidance by moderating horizon-dependent divergence between individual models.

This interpretation is consistent with evidence from large multi-model infectious disease forecasting efforts, which show that forecast accuracy typically decreases with increasing forecast distance, while ensemble-based approaches retain useful early warning information under growing uncertainty (40). More broadly, probabilistic forecasting frameworks emphasise that forecasts are intended to support decisions under uncertainty rather than provide deterministic predictions, particularly at longer forecast ranges (Johansson et al., 2019). An important operational implication is that alerting behaviour can vary across forecast ranges. Because decision thresholds are defined separately for predictions made at different lead times, the number of administrative units flagged as higher risk does not need to remain constant. This flexibility supports decision-making under capacity constraints, allowing higher sensitivity at short lead times while enabling more cautious interpretation of longer-range forecasts. Framing forecast evaluation and use in this way aligns with recent work emphasising the importance of translating forecast metrics into decision-relevant guidance that explicitly accounts for uncertainty, operational capacity, and policy context (41).

### 3.4 Cross-country differences in predictability

Differences in predictive performance across countries reflect how outbreaks are distributed in space, how they tend to spread once they occur, and how consistent these patterns are over time. Across Europe, BTV transmission varies from regions with repeated seasonal circulation to areas where outbreaks occur only sporadically near the limits of suitable environmental conditions. Forecasting approaches developed using outbreak-rich systems such as France, Greece, and Italy may therefore face additional challenges when applied in regions where BTV emergence is rare and historical outbreak data are limited. In practical terms, predictability depends on whether outbreaks tend to appear in many places at once or remain localised, and on how similar outbreak behaviour is across administrative units. Italy represents the most challenging setting because the study period is divided into a large number of administrative units. This increases ecological and operational heterogeneity, as outbreak dynamics differ substantially between provinces. As a result, relationships between predictors and outbreak occurrence that hold in one region may not apply equally well in others. This is reflected in lower average predictive performance and greater variation across regions and prediction windows. In addition, changes in surveillance practices, control measures, or circulating serotypes over time may further reduce the consistency of outbreak patterns, making prediction more difficult (14, 42). In contrast, Greece shows more spatially coherent outbreak patterns at the monthly reporting scale. Once outbreaks occur, they often affect multiple neighbouring areas, leading to similar outbreak timing across regions and stronger persistence over time. Under these conditions, many flexible models are able to identify consistent patterns in the data, resulting in high and relatively uniform predictive performance (43). France lies between these two cases. Using fewer, larger administrative units reduces some local variation and highlights broad seasonal patterns, but outbreaks do not always spread uniformly across the country, limiting overall predictability.

These examples illustrate that the choice of administrative scale strongly influences the way predictable outbreaks appear. Finer spatial divisions preserve local detail but increase variability between units, while larger units combine information across areas, making broad patterns easier to detect but potentially obscuring local differences. In practice, administrative scale is not fixed, and aggregating smaller units into larger ones may reduce local variability and improve predictive stability, although at the cost of reduced spatial resolution. The apparent differences in predictability between countries therefore reflect how outbreak dynamics align with the reporting scale, rather than differences in transmission alone (44). These contrasts also help explain when the logit-weighted ensemble is most useful. In settings where outbreak patterns are broadly similar across regions, such as Greece, many models perform well and the main advantage of the ensemble is providing stable predictions across different prediction horizons. In more heterogeneous settings, particularly Italy and to a lesser extent France, models respond differently to local outbreak patterns as predictions extend further ahead in time. In these cases, combining information across models and adjusting decision thresholds for different forecast ranges produces more consistent performance across regions and time periods.

### 3.5 Relationship to prior forecasting approaches

Prior BTV forecasting has frequently relied on mechanistic or ecological modelling frameworks that explicitly represent temperature dependent vector competence and extrinsic incubation, seasonal *Culicoides* population dynamics, host availability, and spatial spread processes. Within these frameworks, transmission is commonly modelled using thermal performance curves for the extrinsic incubation period, vector survival, and biting rate, often embedded within basic reproduction number or transmission suitability formulations (6, 45).

In mechanistic BTV models, seasonal variation in vector abundance is typically represented using temperature-based or remotely sensed vegetation covariates, rather than being inferred directly from observed outbreak patterns (46, 47), while host density and movement are often incorporated as static or slowly varying modifiers of transmission potential (48, 49). These approaches offer clear biological interpretability and support counterfactual reasoning for intervention planning, including the evaluation of vaccination strategies and vector control under alternative climate or management scenarios. However, their predictive performance can be constrained by parameter uncertainty, strong structural assumptions, and mismatches between idealised transmission processes and the observation mechanisms underlying routine surveillance data. Such limitations become more pronounced under non-stationary conditions, where serotype turnover, intervention changes, or extreme climate events alter the relationship between environmental covariates and reported outbreaks (6, 45, 50–52).

The approach presented here is explicitly predictive and decision oriented. Rather than encoding transmission processes through fixed biological assumptions, it learns flexible mappings from routinely available covariates to future outbreak occurrence and applies horizon specific decision thresholds aligned with surveillance priorities. The strong performance of boosted tree-based learners, together with the additional stability provided by the LWE, indicates that interaction-rich relationships among climate, recent outbreak history, and reporting dynamics can be captured at scale without country-specific redesign or 17 eparameterization. Importantly, this perspective does not position mechanistic and machine learning approaches as competing alternatives. A more practical synthesis is to combine their complementary strengths. Outputs from mechanistic or ecological models, such as temperature-driven suitability indices, estimates of vector abundance, or proxies for transmission potential, can be incorporated as covariates within data driven forecasting ensembles. Conversely, machine learning forecasts can serve as diagnostic tools by flagging departures from expected patterns that may signal regime shifts, changes in surveillance intensity, intervention effects, or violations of mechanistic assumptions. In this way, predictive ensembles and mechanistic models can jointly support both early warning and epidemiological interpretation under changing conditions. The framework relies primarily on routinely available climate summaries and short outbreak-history lags rather than pathogen-specific transmission parameters. Similar forecasting approaches could therefore be applied to other climate-sensitive infectious diseases where comparable surveillance data are available.

### 3.6 Limitations

The outcome analysed here is derived from routine outbreak notifications and therefore reflects a combination of underlying transmission dynamics and the properties of the surveillance process, including incomplete reporting, temporal delays, and changes in case ascertainment over time. As a result, some apparent predictability may arise from stability in reporting behaviour as well as from disease dynamics themselves. In addition, to prioritise portability and comparability across settings, the modelling framework deliberately relies on a restricted predictor set comprising monthly climate summaries and short outbreak-history lags. This choice excludes potentially important but less consistently available determinants of BTV risk, such as vaccination coverage, animal movements, and detailed vector abundance or species composition. These factors are likely to play a larger role at longer forecast horizons and in more heterogeneous epidemiological settings, and their omission defines the scope within which the present results should be interpreted. The modelling framework does not explicitly represent *Culicoides* species composition or vector ecological heterogeneity and therefore cannot disentangle species-specific transmission mechanisms across regions. In addition, other drivers not explicitly represented here include variation among circulating BTV serotypes, broader environmental changes, and socio-economic factors that influence livestock movement and connectivity, such as trade networks, production systems, and management practices, all of which may affect vector abundance and outbreak risk. Differences in surveillance intensity, reporting practices, administrative granularity, and data availability across countries may influence apparent predictability and limit direct comparability, particularly in regions with sparse or inconsistent surveillance data. These disparities highlight the need for improved surveillance harmonisation to support more proactive forecasting, especially in regions where BTV outbreaks are currently under-reported or poorly characterised. In addition, because the modelling framework relies partly on previous outbreak observations, its predictive utility may be limited in areas with little or no historical outbreak data, where preparedness applications would require additional ecological or mechanistic information.

### 3.7 Future directions

Future work could enhance both forecast skill and operational relevance by incorporating additional predictors that remain broadly available but better capture local transmission conditions. These include higher-resolution remotely sensed climate and land-surface variables, vegetation and soil-moisture proxies linked to vector habitat suitability, and routinely collected indicators of vaccination coverage, livestock density, or entomological surveillance where available. Beyond predictor expansion, explicitly representing spatial dependence and spread processes, for example through spatial lags or movement-informed connectivity features, may further improve performance in settings with strong spatial coupling. From an operational perspective, prospective deployment should prioritise external validation, alongside ongoing monitoring of calibration, alert rates, and horizon-specific performance, enabling drift-aware recalibration and adaptive updating of models or decision thresholds as epidemiological and surveillance conditions evolve. Extending proactive outbreak forecasting to regions with limited or no historical outbreak data remains a key challenge. In such settings, reliance on outbreak history lags is insufficient, and additional information sources, such as entomological surveillance, livestock movement data, vaccination coverage, or mechanistic suitability indices, will be required to support early warning at appropriate spatial granularity. Future work could also benefit from greater integration and harmonisation of surveillance data across countries. While the present analysis reflects the structure of country-specific reporting systems, BTV transmission does not respect administrative borders. Incorporating cross-border risk indicators, regional climate summaries, or spatial lags may further improve predictive performance, particularly in border regions and transboundary risk corridors.

## 4. Conclusion

This study demonstrates that a compact LWE can provide reliable multi-horizon early warning forecasts for BTV outbreaks using routinely available climate variables and lagged outbreak history. Across France, Italy, and Greece, predictive skill decreases with lead time, but the LWE remains consistently strong under the recall-oriented *𝐹*_2_ objective and becomes the top-ranked method at longer horizons (Month +3 to Month +6), where single-model rankings are least stable. This indicates that the main contribution of the LWE is not only high average performance, but greater robustness to horizon-dependent instability under heterogeneous spatio-temporal conditions --- an essential property for surveillance settings in which missed outbreaks carry higher costs than manageable false alarms. Operationally, informative risk estimates up to six months ahead can support earlier and more proportionate responses. Short-horizon forecasts can guide targeted surveillance escalation and diagnostic prioritisation, while intermediate and longer horizons can inform resource planning, vaccination logistics, and preparedness actions that are subsequently refined as new data arrive. More broadly, the proposed horizon-aware ensemble framework provides a scalable foundation for forecasting other climate-sensitive, transboundary vector-borne diseases, and contributes a practical component for One Health preparedness as environmental conditions and pathogen ranges continue to shift. Overall, these results show that robust, horizon-aware ensemble forecasting using routinely available data can deliver transferable and decision-relevant early warning signals for vector-borne disease surveillance under heterogeneous and changing conditions.

## 5. Methods

### 5.1 Study design and data

#### 5.1.1 Study areas

Unit–month BTV occurrence datasets were assembled for three European settings, France, Greece, and Italy, to evaluate whether a single multi-horizon forecasting framework can generalise across heterogeneous epidemiological, spatial, and climatic contexts. These countries were selected due to their history of repeated BTV outbreaks and because they fall within the highest category of reported BTV records in Europe over the study period (Figure 2)(53–55). The datasets are defined based on official veterinary reporting units, which differ in operational reporting scale across countries: France uses coarser spatial aggregation, whereas Greece and Italy are subdivided into finer units (Figure 2B). Established surveillance systems and access to comprehensive climate and epidemiological data in all three countries provide a strong foundation for comparative spatiotemporal analysis of outbreak patterns (53–56). Table 2 summarises the number of reporting units and the resulting unit–month outcome prevalence in the development period (2005–2021), used for model training and internal validation, and the test period (2022–2024), used to evaluate predictive performance on future observations.

**Figure 2:**
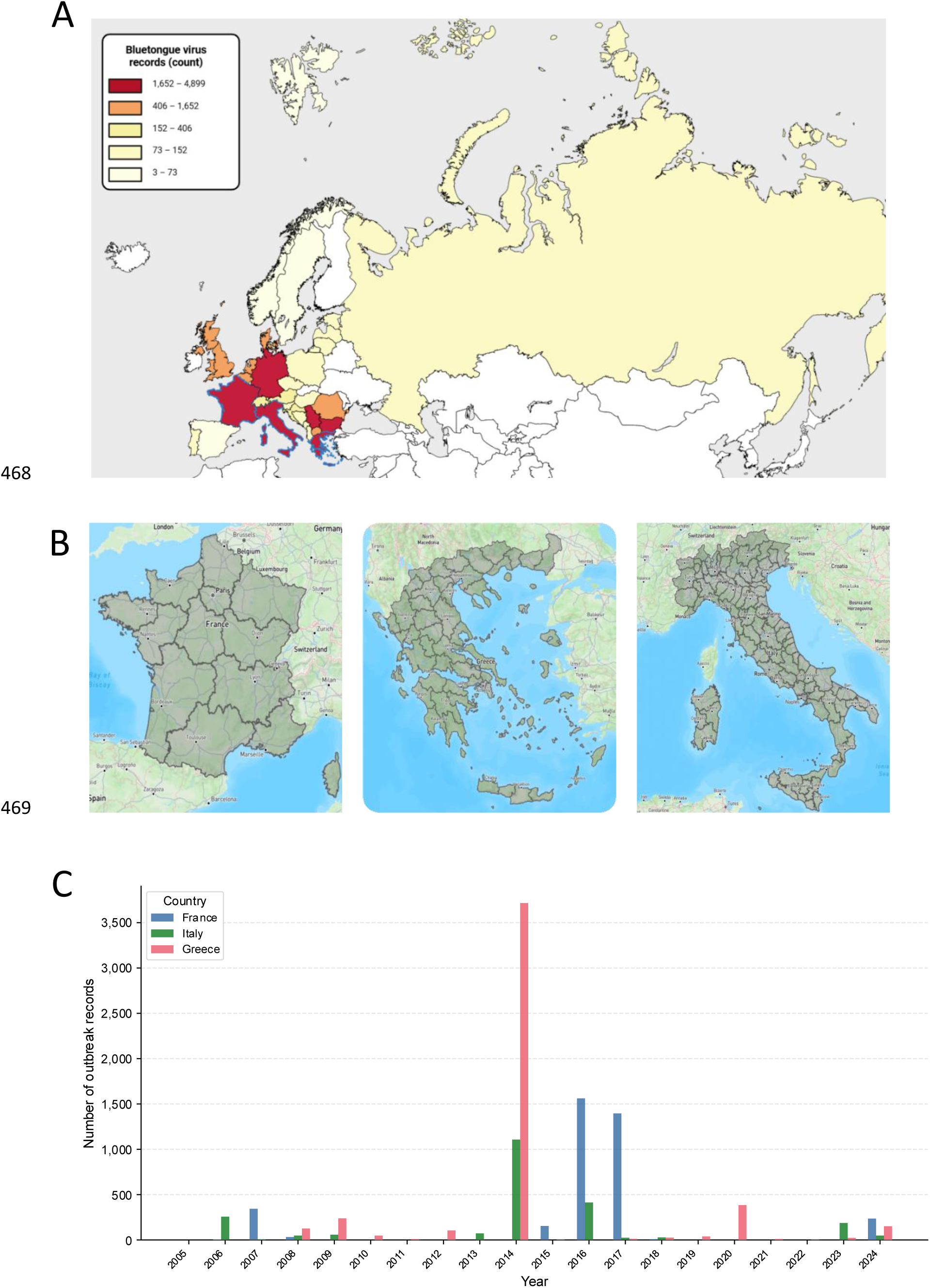
Study context and spatial units used for multi-horizon bluetongue virus (BTV) forecasting**. (A)** European map of reported BTV outbreak records over the study period, aggregated at the country level. Countries are shaded by total numbers of reported outbreaks using a sequential colour scale from pale yellow through orange to dark red, with darker colours indicating higher record counts. France, Italy, and Greece fall within the highest outbreak category and are highlighted as the study settings with blue country borders. **(B)** Study areas and administrative reporting units used for multi-horizon BTV forecasting in France, Italy, and Greece. Shaded polygons show the spatial units at which outbreak occurrence is defined and predicted at a monthly reporting scale. Forecasts are generated separately for each unit–month combination using covariates aggregated to this administrative level. Each reporting unit is linked to gridded climate variables and historical outbreak records and is represented by a single reference coordinate for climate data extraction and model indexing. **(C)** Annual number of reported BTV outbreak records in France, Italy, and Greece over the study period. Bars show the total number of outbreak records per year for each country.

**Table 2:** Summary of the modelling dataset and outcome prevalence at the administrative unit–month scale for each study country. Each observation represents one administrative reporting unit in one calendar month. A unit–month is classified as positive if at least one confirmed bluetongue virus (BTV) outbreak is reported in that unit during the month. The table reports the number of reporting units, the total number of months analysed, the resulting number of unit–month observations, and the number and proportion of positive unit–months. Prevalence during the model development period (January 2005–December 2021) and during the held-out evaluation period (January 2022–December 2024) are shown separately to illustrate differences between training and test data.

| Country | Units (U) | Months (T) | Unit-months | Positive | Prevalence (%) | Train prevalence (%) | Test prevalence (%) |
| --- | --- | --- | --- | --- | --- | --- | --- |
| France | 13 | 240 | 3,900 | 1,060 | 27.18 | 29.49 | 59.4 |
| Greece | 55 | 240 | 16,500 | 2,292 | 13.89 | 8.57 | 67.1 |
| Italy | 107 | 240 | 32,100 | 1,316 | 4.10 | 5.59 | 2.4 |

France spans both Mediterranean and north-western European climatic zones and has experienced repeated BTV incursions during recent expansion waves in Europe (53, 57). BTV activity in France has shifted from historically sporadic detection to more recurrent circulation, including re-emergence of BTV-8 following earlier elimination (53, 57, 58). Transmission occurs within temperate European vector systems, allowing outbreaks to reappear with the resumption of seasonal vector activity.

Greece lies predominantly within the Mediterranean climatic zone and comprises a mixture of mainland and island regions. BTV outbreaks in Greece have been characterised by episodic but extensive transmission events, including the 2014 epidemic during which cases spread rapidly across mainland and island regions within a single vector season (59–61). Continued surveillance indicates ongoing BTV activity consistent with a system where warm-season vector suitability enables rapid national-scale dissemination once transmission becomes established (61).

Italy also lies largely within the Mediterranean climatic zone and has experienced repeated BTV circulation in several regions, particularly in southern and insular areas (62). BTV transmission in Italy is considered established in specific regions and typically occurs on a seasonal basis. Evidence from Mediterranean island systems such as Sardinia suggests that extended seasonal vector activity may facilitate persistence of BTV between transmission seasons (62).

#### 5.1.2 Outbreak data

Records of BTV outbreaks between January 2005 and December 2024 were obtained from the World Animal Health Information System (WAHIS), maintained by the World Organisation for Animal Health (WOAH) (63). Outbreaks recorded in WAHIS are officially reported by national veterinary authorities and include information on outbreak timing, affected host species, and geographic location. For this study, the geographic coordinates of each reported outbreak were extracted and assigned to official veterinary reporting units using point-in-polygon matching. Outbreaks were then aggregated at the unit–month level to define a binary outcome ≥ *𝑦_𝑢,𝑡_*, indicating the presence of at least one confirmed outbreak in unit *𝑢* during month *𝑡* (*𝑦_𝑢,𝑡_* = 1 if 1 outbreak occurred, and *𝑦_𝑢,𝑡_* = 0 otherwise). In this study, outbreak occurrence refers to the presence of at least one officially reported outbreak notification within an administrative unit during a given month, as recorded in WAHIS surveillance data. This formulation reflects an early warning triage perspective, in which the primary question is whether transmission is occurring in a given unit–month. Where this information was available in WAHIS records, suspected or unconfirmed notifications were excluded. Outbreaks were aggregated across host species to reflect unit-level transmission reporting rather than host-specific incidence. The distribution of outbreak records by host species across the three study countries is summarised in Table 3. The majority of outbreak records involve cattle, sheep, and goats, although the relative composition varies across countries, with cattle dominating records in France and small ruminants more frequently reported in Greece and Italy, while other species were recorded only sporadically.

**Table 3.** Distribution of reported BTV outbreak records by host species and country in the modelling dataset (2005–2024). Outbreak records were obtained from the World Animal Health Information System (WAHIS) maintained by the World Organisation for Animal Health (WOAH). Counts represent the number of confirmed outbreak notifications attributed to each host species within each country prior to aggregation for modelling. For the forecasting framework, outbreak records were subsequently aggregated across host species at the administrative unit–month level to define binary unit-level transmission events. Cattle, sheep, and goats account for the majority of records, while other species were reported only sporadically.

| Species | France | Greece | Italy |
| --- | --- | --- | --- |
| Cattle | 3498 | 153 | 788 |
| Sheep | 231 | 3372 | 1322 |
| Goats | 9 | 1182 | 127 |
| Sheep/goats (mixed herd) | 2 | 191 | 0 |
| Buffalo | 0 | 0 | 7 |
| Wild sheep | 0 | 0 | 3 |
| Yak | 0 | 0 | 1 |
| Camelids | 0 | 0 | 1 |
| Fallow deer | 0 | 0 | 1 |
| Alpaca | 0 | 0 | 1 |

#### 5.1.3 Climate data

Climatic predictors were derived from the E-OBS gridded observational dataset distributed via the Copernicus Climate Data Store (64). Monthly gridded climate variables covering the period 2005–2024 were extracted at a spatial resolution of 0.1^◦^ (approximately 10km). Variables were selected based on established links between meteorological conditions and *Culicoides* survival, activity, dispersal, and temperature-dependent virus replication, which together shape BTV transmission potential (7, 65). The climate feature set included mean, minimum, and maximum 2m air temperature, total precipitation, relative humidity, surface pressure, and 10m wind speed. Gridded values were aggregated to the reporting-unit level by averaging across all grid cells intersecting each unit polygon; precipitation was summed within each month to represent cumulative moisture availability and wet-season carryover (66).

#### 5.1.4 Additional predictors

To better capture short-term dynamics relevant to vector-borne transmission, several derived predictors were constructed from the monthly climate variables. Monthly temperature variability was represented using the temperature range, defined as 𝑇_𝑢,𝑡_^max^−𝑇_𝑢,𝑡_^min^. Recent moisture conditions were summarised using a rolling 6-month precipitation total,

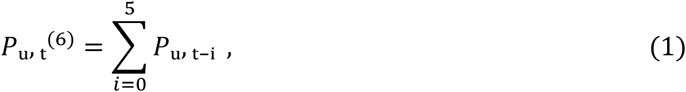

computed from monthly precipitation totals. To account for recent local persistence and temporal dependence that are operationally informative in surveillance and early warning contexts, outbreak-history lags were included as six binary features, {𝑦_𝑢,𝑡−1_,…,𝑦_𝑢,𝑡−6_ (40). These lagged indicators are interpreted as predictive signals of ongoing transmission rather than as causal drivers. Table 4 summarises all predictors, along with their units, temporal aggregation, and data sources.

**Table 4:** Predictor variables used for multi-horizon BTV outbreak forecasting, with units, temporal aggregation, and data sources Climate variables are derived from gridded E-OBS and Copernicus datasets and aggregated to monthly summaries at the administrative reporting-unit level. Six-month rolling precipitation corresponds to the cumulative precipitation over the preceding six months. Outbreak history lags represent binary indicators of reported outbreak presence in the same reporting unit during each of the previous one to six months. The target outcome is a binary indicator of outbreak presence in a given unit and month, defined as at least one confirmed BTV outbreak record reported during that month. All outbreak-derived variables are constructed from aggregated WAHIS surveillance.

| Feature | Units | Temporal aggregation | Source |
| --- | --- | --- | --- |
| Mean air temperature at 2 m | °C | Monthly mean | E-OBS / Copernicus |
| Minimum air temperature at 2 m | °C | Monthly minimum | E-OBS / Copernicus |
| Maximum air temperature at 2 m | °C | Monthly maximum | E-OBS / Copernicus |
| Air temperature range | °C | Monthly mean | Derived from E-OBS |
| Total precipitation | mm | Monthly total | E-OBS / Copernicus |
| Six-month rolling precipitation | mm | Monthly total | Derived from E-OBS |
| Relative humidity | % | Monthly mean | E-OBS / Copernicus |
| Surface pressure | hPa | Monthly mean | E-OBS / Copernicus |
| Wind speed at 10 m | m s<sup>-1</sup> | Monthly mean | E-OBS / Copernicus |
| Outbreak history lags (previous 1–6 months) | Binary (0/1) | Monthly | Derived from WAHIS |
| Latitude and longitude | degrees | Static | Processed dataset |
| Outbreak presence (target variable) | Binary (0/1) | Monthly | Aggregated WAHIS |

### 5.2 Prediction framework and data structuring

#### 5.2.1 Data harmonisation and completeness

Outbreak notifications and climate covariates were organised into complete unit-specific monthly time series, aligning all predictors by calendar month across datasets. Months with no confirmed outbreaks were treated as true non-events (*𝑦_𝑢,𝑡_* = 0). Climate covariates were spatially aggregated to reporting-unit polygons and used to construct derived predictors. The resulting unit–month table contained complete predictor coverage for all included observations, and therefore no imputation was required.

#### 5.2.2 Sliding-window formulation and targets

Forecasting was framed as a multi-horizon binary classification task (Figure 3). For each unit *𝑢* and month *𝑡*, the model uses the most recent six months of predictors (*𝐿* = 6) as input *𝑥_𝑢,𝑡_*. The aim is to predict whether an outbreak will be reported *ℎ* months ahead, for horizons *ℎ* = 1,…,6, using targets *𝑦_𝑢,𝑡_*_+_*_ℎ_* (equal to 1 if an outbreak occurs in month *𝑡* + *ℎ*, and 0 otherwise). This formulation captures short-term temporal dependence and seasonal accumulation of outbreak risk while maintaining a compact input structure. Forecast horizons of up to six months therefore span both near-and medium-term prediction windows across diverse epidemiological settings.

**Figure 3:**
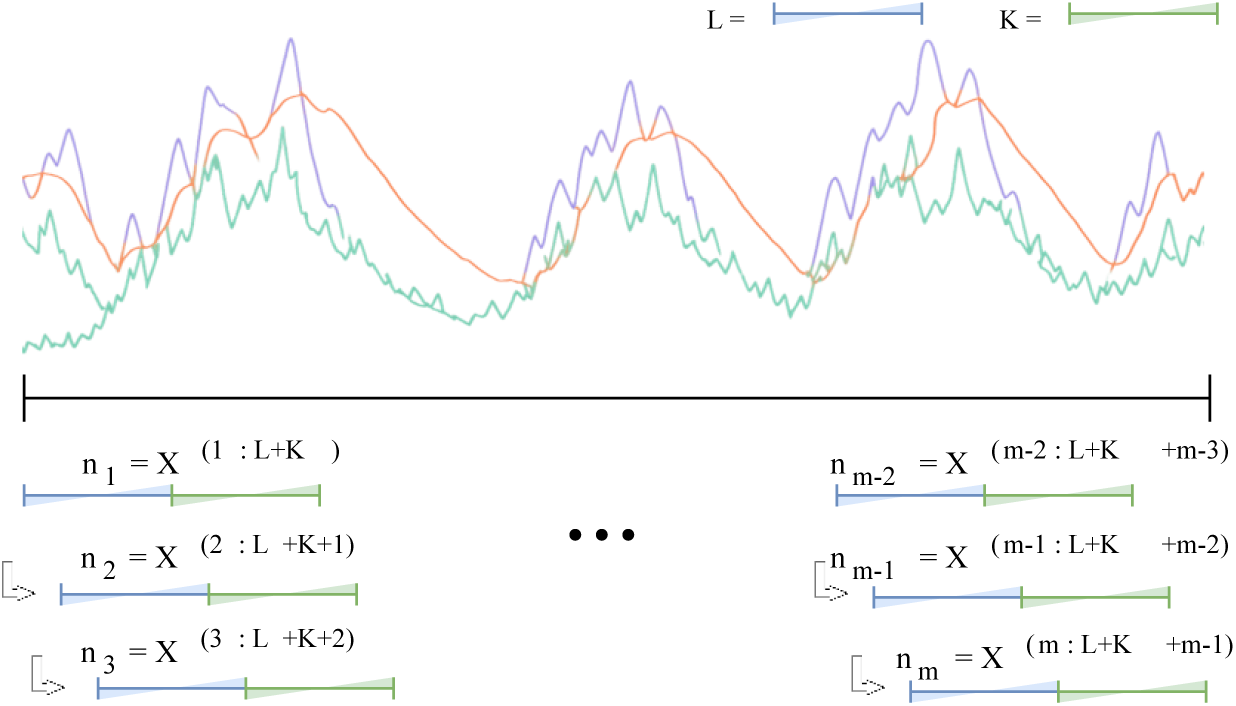
Sliding-window construction for multi-horizon outbreak forecasting at the unit–month scale. Coloured curves illustrate an example monthly time series of predictor variables for a single administrative reporting unit. For a given reference month *𝑡*, the model input consists of the most recent *𝐿* =6 months of predictors (blue window). Binary outcome targets are defined at forecast horizons *ℎ* =1,…,6 months ahead (green offset), indicating whether at least one BTV outbreak is reported in the same unit at month *𝑡* + *ℎ*. As the window advances by one month, overlapping input–target pairs are generated along the time series, producing a sequence of supervised samples that capture short-term dependence and seasonal build-up while supporting near-to medium-term forecasts.

### 5.3 Baseline models

A diverse set of supervised classifiers was benchmarked to assess how different inductive biases perform under nonlinear relationships and severe class imbalance. The model suite includes interpretable linear baselines, distance and margin-based nonlinear methods, and tree-based ensemble learners commonly used for tabular prediction. Tree ensemble methods (especially gradient-boosted trees) are widely recommended as strong baselines for real-world tabular prediction tasks (67).

Models evaluated in this study included regularised, class-weighted logistic regression (68); *k*-Nearest Neighbours (69); linear and RBF-kernel support vector classifiers (70); Random Forest (71) and Extra Trees (72); Balanced Random Forest (73); and boosting-based ensembles including Gradient Boosting (74), AdaBoost (75), HistGradientBoosting, XGBoost (76), LightGBM (77), and CatBoost (78). For algorithms that support it, class-balanced loss weighting was applied to mitigate class imbalance; Balanced Random Forest uses class-balanced bootstraps by design. Models that do not directly output calibrated probabilities suitable for threshold-based decision-making (e.g., Linear SVM and RidgeClassifier) instead produce decision scores that are not directly comparable across methods or forecasting horizons. For these models, decision scores were converted to probabilities using sigmoid (Platt) calibration fitted on the development data (Platt et al., 1999). Calibration was performed within the training pipeline to avoid leakage from the held-out test period.

### 5.4 Logit-weighted ensemble method

#### 5.4.1 Motivation and overview

Single-model forecasting has limitations in this setting for two reasons. First, predictive performance degrades with increasing lead time, such that modest differences in calibration and operating thresholds under class imbalance can result in substantial divergence in recall at longer horizons (Month +3 to Month +6). Second, heterogeneity across administrative units and countries implies that different learners perform best under different data regimes (e.g., near separable versus noisy settings), leading to horizon-dependent rank reversals and instability in alert generation. To mitigate these failure modes, a LWE framework is employed, pooling complementary decision functions from multiple strong learners to reduce sensitivity to overconfident probability estimates and to enable horizon-specific operating point selection. The LWE is constructed on the development data in four phases: logit transformation, top-*𝑘* model selection, learned logit-weighted aggregation, and horizon-specific thresholding.

#### 5.4.2 Logit transformation

Predicted outbreak probabilities *𝑝^_𝑚,ℎ_*(*𝑥*) are obtained from each base model *𝑚* at forecast horizon *ℎ*. Aggregation on the probability scale is sensitive to extreme predictions near 0 or 1, a behaviour that is common under severe class imbalance. To stabilise aggregation and obtain an additive representation, predicted probabilities are transformed to the log-odds scale and clipped to [*𝜖,*1 − *𝜖*] to ensure numerical stability. The logit-transformed scores are defined as

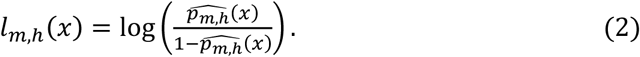

#### 5.4.3 Top-𝒌 ensemble membership

Candidate base learners are filtered to construct a compact ensemble representation. Base models are ranked according to mean validation *𝐹*_2_, averaged across forecasting horizons using horizon-specific thresholds, and the top-*𝑘* models are retained. This filtering step avoids dilution of ensemble performance by excluding poorly calibrated or unstable learners at longer horizons, while maintaining diversity among consistently competitive models. The choice of ensemble size *𝑘* was guided by validation performance, with the mean *𝐹*_2_ score evaluated across candidate values of *𝑘* (Figure 4).

**Figure 4:**
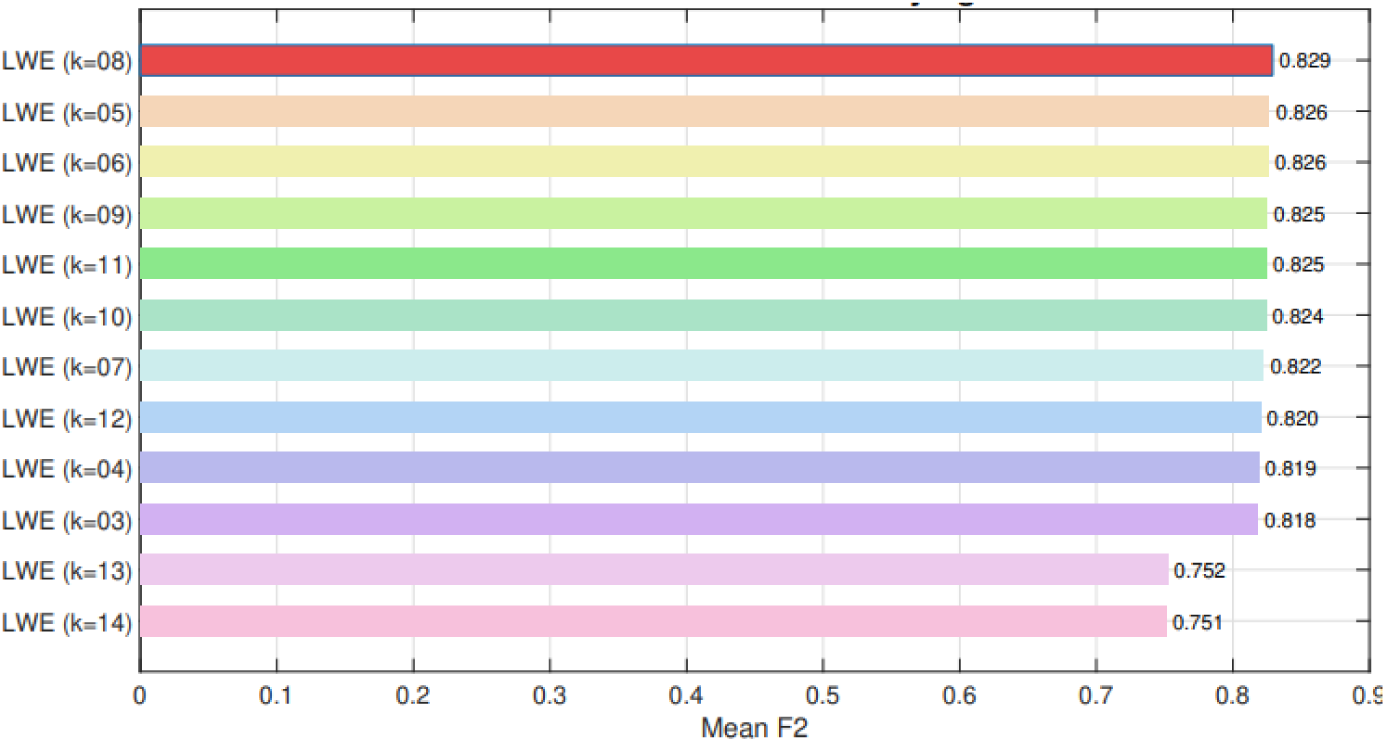
Logit-weighted ensemble (LWE) size selection on the development/validation split. Mean 𝐹_2_ score as a function of ensemble size 𝑘, obtained by combining the top-𝑘 base models ranked by validation performance. The ensemble size 𝑘 = 8 was selected using validation data only and fixed prior to evaluation on the held-out test period; results across alternative values of 𝑘 are shown for transparency.

#### 5.4.4 Learned logit-weighted aggregation

Base learners can be informative in different regimes (e.g., across countries, administrative units, or forecast horizons), making fixed or uniform averaging suboptimal. For each forecast horizon *ℎ*, non-negative weights are optimised on the validation split to emphasise models that are most informative at that lead time. Let 𝑀_𝑘_denote the selected set of base models, with weights 𝑤_𝑚,ℎ_ ≥ 0 constrained such that ∑_𝑚∈ℳ𝓀_ 𝑤_𝑚,ℎ_ = 1. _Logit-scale predictions_ are aggregated as

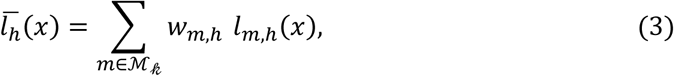

and mapped back to the probability scale using the logistic function,

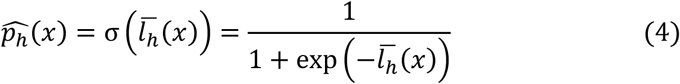

#### 5.4.5 Horizon-specific thresholding

Producing actionable binary outputs requires the specification of a decision threshold; however, the optimal operating point varies with forecast horizon because outcome prevalence and score separability change with lead time. A horizon-specific threshold *𝜏_ℎ_* is therefore selected on the validation data to maximise the *𝐹*_2_ score over a grid of candidate thresholds (0.05–0.95). Final predictions are defined as

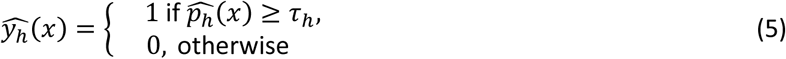

### 5.5 Evaluation design and implementation

#### 5.5.1 Time-respecting design

A strict time-based data split was applied. The development period spans January 2005 to December 2021, and the held-out test period spans January 2022 to December 2024. A sliding-window sample with reference month *𝑡* and horizon *ℎ* was assigned to the test set if its target month *𝑡* + *ℎ* fell within the 2022–2024 period; otherwise, it was assigned to the development set, preventing information leakage from windows whose look-back period overlaps the test era. Within each country, the development data were further partitioned in a time-respecting manner, with the final 15% of months reserved for validation.

#### 5.5.2 Training and prediction procedure

Horizon-specific classifiers were trained separately for each forecast horizon (*ℎ* = 1,…,6). For scale-sensitive algorithms (logistic regression, *𝑘*-nearest neighbours, and support vector machines), predictors were standardised using StandardScaler (79), with scaling parameters estimated on the training data and applied to subsequent data splits through a pipeline. Hyperparameters were selected on the development data using validation *𝐹*_2_ (80); where applicable, probability calibration and horizon-specific decision thresholds were also determined using development data only. Final probabilistic forecasts and corresponding binary predictions (after thresholding) were generated for the held-out test period (2022–2024).

#### 5.5.3 Implementation

All models were implemented in Python 3.11 using scikit-learn 1.5 (Pedregosa et al., 2011), XGBoost 2.1, CatBoost 1.2.5, and LightGBM 4.6.0.

#### 5.5.4 Performance metrics

Models are evaluated under a recall-prioritised outbreak-detection objective, reflecting that failing to flag an outbreak month (a false negative) is typically more consequential than issuing a manageable false alarm (a false positive). Accordingly, the primary performance metric is the *𝐹*_2_ score, a member of the *𝐹_𝛽_* family that explicitly upweights recall relative to precision (81). Let TP, FP, and FN denote the numbers of true positives, false positives, and false negatives, respectively. Precision measures the reliability of positive predictions, while recall measures how completely outbreaks are detected:

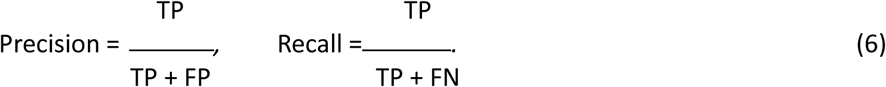

The *𝐹_𝛽_* score combines these quantities via a weighted harmonic mean,

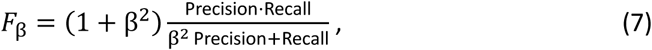

where *𝛽 >* 1 places greater emphasis on recall than precision (and *𝛽 <* 1 does the opposite). The *𝐹*_2_ score is obtained by setting *𝛽* = 2, which weights recall four times as strongly as precision

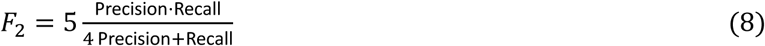

For each forecast horizon *ℎ*, threshold-dependent metrics (including *𝐹*_2_) are computed using horizon-specific decision thresholds selected on the validation split. Unless stated otherwise, overall performance across horizons is summarised as the arithmetic mean of horizon-specific scores.

#### 5.5.5 Pooled evaluation setup

Benchmark models were evaluated on a pooled test set combining unit–month windows from France, Italy, and Greece. Pooled performance is reported as a micro-average by concatenating all test predictions across countries, weighting each country by its number of test windows. To avoid masking heterogeneity, pooled summaries are reported alongside country-specific results (Table 1).

## CRediT authorship contribution statement

Lucy M. Devlin: Conceptualisation; Data curation; Formal analysis; Investigation; Methodology; Project administration; Resources; Software; Validation; Visualisation; Writing – original draft; Writing – review & editing. Phi H. Nguyen: Data curation; Formal analysis; Investigation; Methodology; Software; Validation; Visualisation; Writing – original draft; Writing – review & editing. Son T. Mai: Conceptualisation; Methodology; Formal analysis; Project administration; Supervision; Writing – review & editing. Ross N. Cuthbert: Conceptualisation; Project administration; Supervision; Writing – review & editing. Phu Ngoc Doan: Data curation; Formal analysis; Software. Viet Hung Tran: Data curation; Formal analysis; Software. Zichi Zhang: Data curation; Formal analysis; Software. Archie K. Murchie: Supervision; Writing – review & editing. Connor G. G. Bamford: Supervision; Writing – review & editing. Jaimie T. A. Dick: Supervision; Writing – review & editing. Eric R. Morgan: Supervision; Writing – review & editing.

## Declaration of competing interest

The authors declare that they have no known competing financial interests or personal relationships that could have appeared to influence the work reported in this paper.

## Acknowledgements

This work was funded by the Department of Agriculture, Environment and Rural Affairs (DAERA). The authors gratefully acknowledge this support.

## Data and code availability

All data used in this study are publicly available from the original sources cited in the manuscript.

## Notes

### Competing Interest Statement

The authors have declared no competing interest.

